## Supplementary Figure S1 for "Spatial periodicity in grid cell firing is explained by a neural sequence code of 2-D trajectories"

### SUPPLEMENTARY INFORMATION

#### Supplementary Figure S1

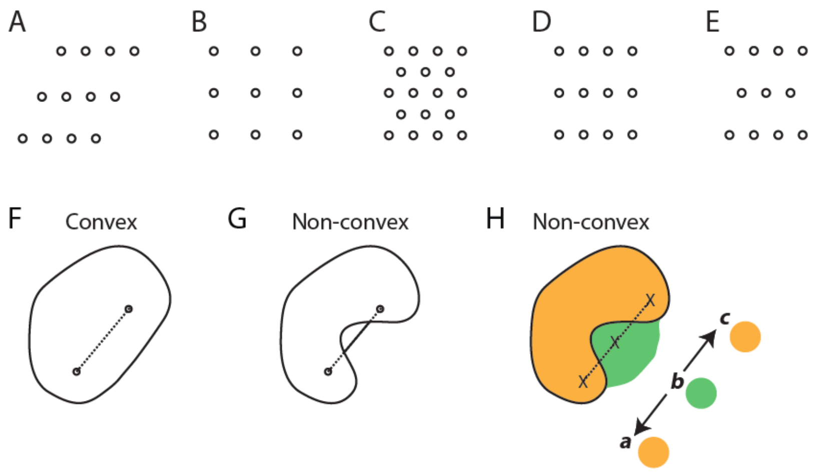


**Supplementary Figure S1. Trajectory coding by cell sequences requires a dense packing of convex firing fields. A-E,** Examples of the five types of lattices in the plane. A, Oblique. B, Square. C, Hexagonal. D, Rectangular. E, Centered rectangular. **F,G,** Examples of a convex and non-convex geometric object such as a possible spatial firing field of a neuron. **H,** A non-convex spatial firing field that results in an ambiguous cell sequence code for trajectories in space. In this example, two spatial firing fields of two cells are shown, indicated by green and orange colors. The three crosses mark three locations in space. The sequence *green* → *orange* can represent two opposite trajectories, namely *b* → *a* and *b* → *c*.
