## Supplementary Figure S2 for "Spatial periodicity in grid cell firing is explained by a neural sequence code of 2-D trajectories"

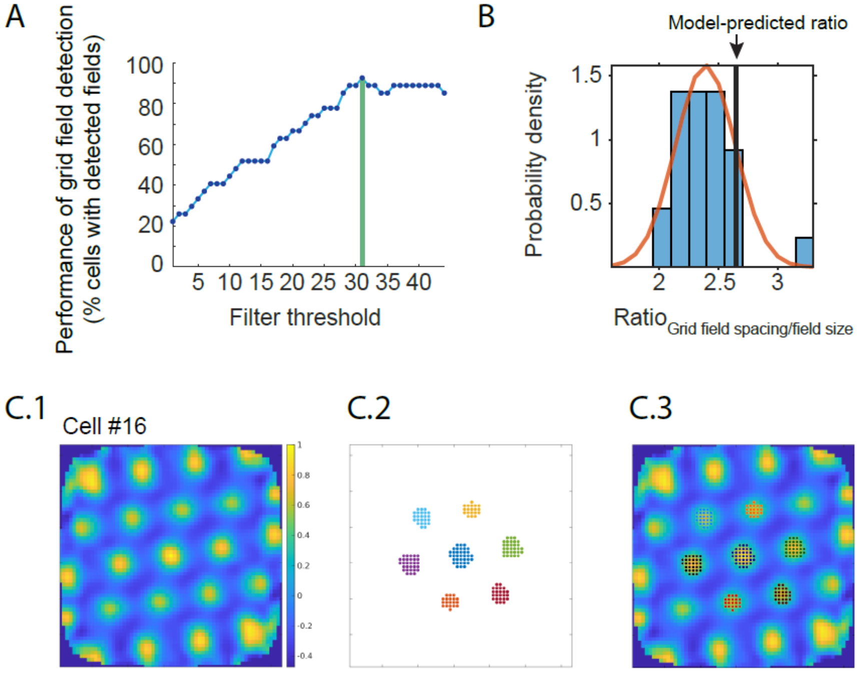


**Supplementary Figure S2. Quantification of grid field spacing and grid field size in grid cells obtained from mice.** We analyzed a total of 27 grid cells from a previously published study[42] to quantify the ratio between grid field spacing and grid field size. The experimental data on grid cells were obtained from mice freely foraging in a 1 x 1 $m^2$ square environment with walls and a visual cue card during baseline recording sessions. The model-predicted value of the ratio of grid field spacing to a diameter-like metric of field size is $\sqrt{7} \approx 2.65$. Note that this reflects the ideal ratio assuming no noise in experimental measurements, no transient drifts of grid maps, no path integration error, no conjunctive coding or any other factors that could result in out-of-field firing of recorded neurons. The grid field size measured from firing rate maps of experimental data is, therefore, expected to be larger than the model-predicted grid field size, resulting in smaller ratios of grid field spacing/field size. **A,** Successful identification of grid fields for quantification of grid field size is a function of the applied filter threshold (see Methods). The algorithm performed best (measured as the percentage of cells where grid fields could be detected) with a filter threshold of 31% of the peak firing rate (vertical green line). **B,** Histogram of the ratios measured in n = 25 grid cells out of 27 grid cells, obtained with applying the 31% filter threshold. As expected, the model-predicted value is larger but falls within the distribution of experimental data with mean = 2.44 and standard deviation = 0.23. The red line shows a normal distribution with the experimentally observed mean and standard deviation. **C,** Performance of the grid field detection algorithm illustrated for one example neuron (cell #16). **C.1,** The spatial autocorrelogram of the firing rate map of cell #16. **C.2,** Grid fields detected by the field detection algorithm with the filter threshold set to 31% of the peak firing rate. Colors identify individual grid fields. The measured ratio (grid field spacing/field size) of this cell was 2.48. **C.3**, Detected grid fields merged with spatial autocorrelogram.
