## Supplementary Figure S3 for "Spatial periodicity in grid cell firing is explained by a neural sequence code of 2-D trajectories"

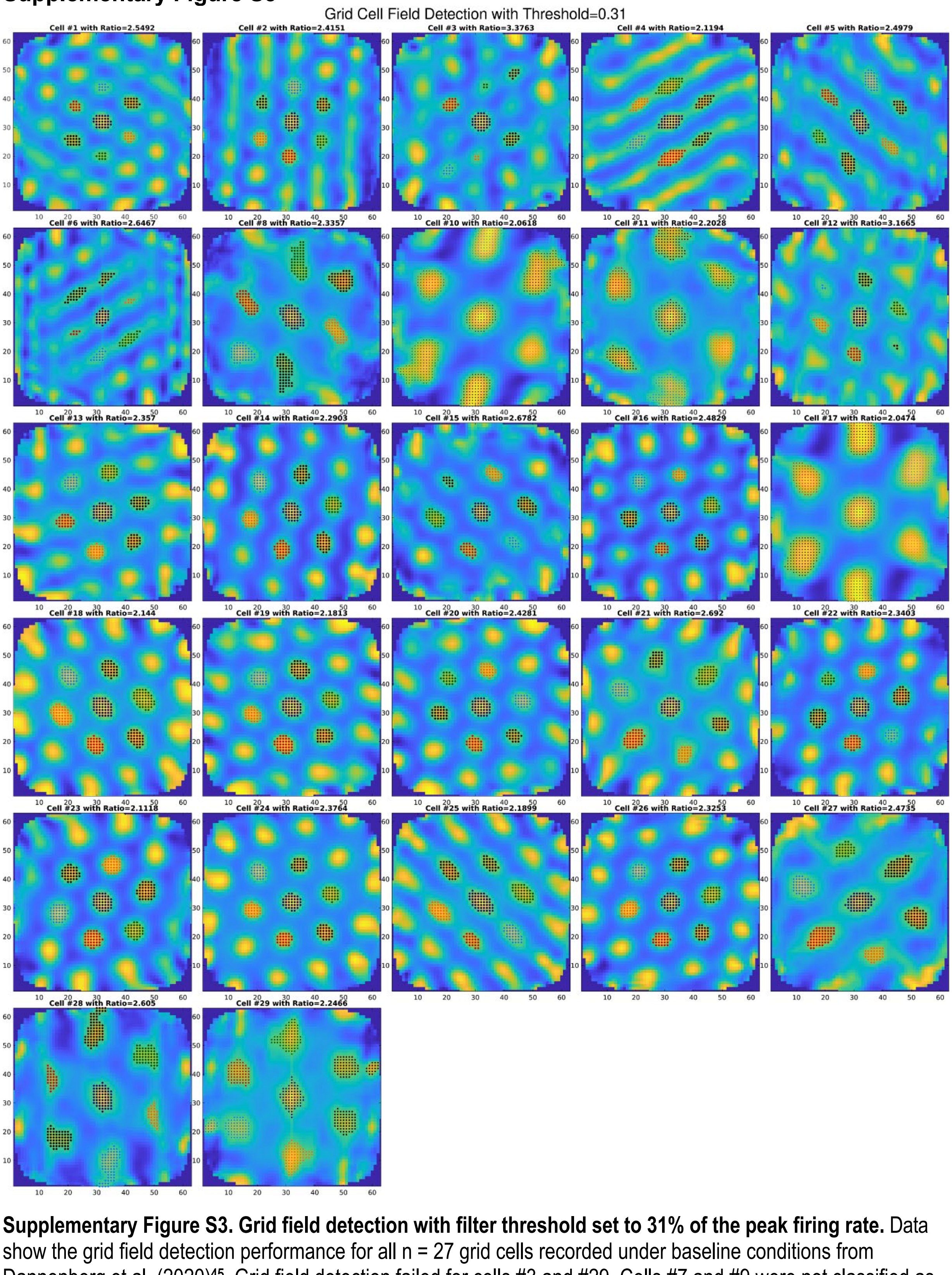


**Supplementary Figure S3. Grid field detection with filter threshold set to 31% of the peak firing rate.** Data show the grid field detection performance for all n = 27 grid cells recorded under baseline conditions from Dannenberg et al. (2020)[45]. Grid field detection failed for cells #3 and #29. Cells #7 and #9 were not classified as grid cells during baseline recording sessions and, therefore, excluded for this analysis.
