## Supplementary Figure S4 for "Spatial periodicity in grid cell firing is explained by a neural sequence code of 2-D trajectories"

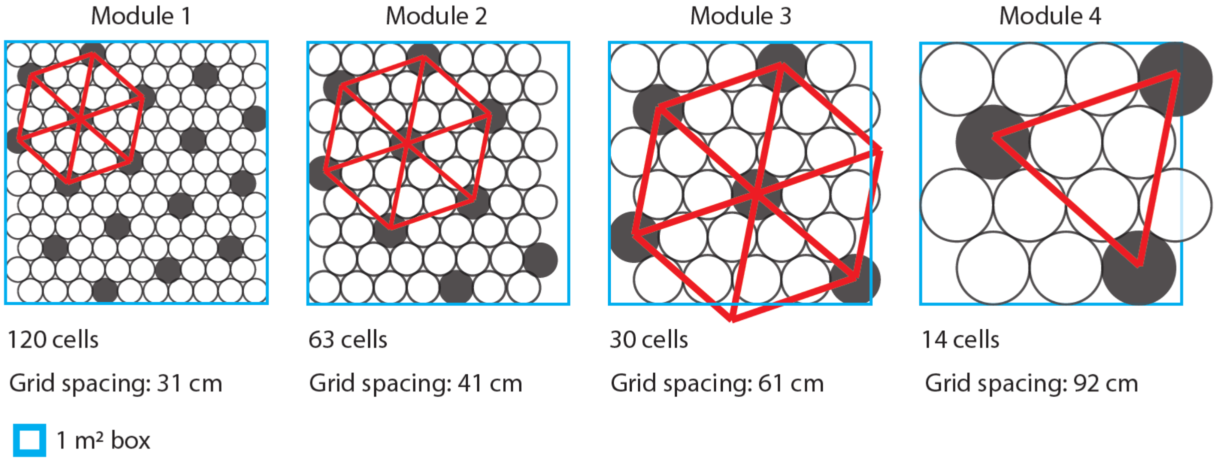


**Supplementary Figure S4. Grid cell modules provide a pyramidal parametric sampling of 2-D space enabling simultaneous multiscale representation of space.** The sampling of 2-D space by grid fields is cut in half from module to module so that each grid cell module represents space at half of the resolution of the previous module. This resembles Gaussian pyramids or mip maps in computer vision. Representation of 2-D space by different grid cell modules thereby allows the simultaneous representation of space at meaningful spatial resolutions in a computationally efficient way. The blue box represents a 1-m x 1-m square environment. Circles represent densely packed grid fields from multiple grid cells. Filled circles represent grid fields of a single grid cell. Grid fields are enlarged in space from module to module so that the total number of grid fields sampling the space approximately doubles from module to module. While the total number of grid fields doubles from module to module, the spacing between the grid fields of an individual grid cell increases by a factor of $\sqrt{2}$.
